# Image-derived Models of Cell Organization Changes During Differentiation of PC12 Cells

**DOI:** 10.1101/522763

**Authors:** Xiongtao Ruan, Gregory R. Johnson, Iris Bierschenk, Roland Nitschke, Melanie Boerries, Hauke Busch, Robert F. Murphy

## Abstract

Cellular differentiation is a complex process requiring the coordination of many cellular components. PC12 cells are a popular model system to study changes driving and accompanying neuronal differentiation. While significant attention has been paid to changes in transcriptional regulation and protein signaling, much less is known about the changes in cell organization that accompany PC12 differentiation. Fluorescence microscopy can provide extensive information about this, although photobleaching and phototoxicity frequently limit the ability to continuously observe changes in single cells over the many days that differentiation occurs. Here we describe a generative model of differentiation-associated changes in cell and nuclear shape and their relationship to mitochondrial distribution constructed from images of different cells at discrete time points. We show that our spherical harmonic-based model can accurately represent cell and nuclear shapes by measuring reconstruction errors. We then learn a regression model that relates cell and nuclear shape and mitochondrial distribution and observe that the predictive accuracy generally increases during differentiation. Most importantly, we propose a method, based on cell matching and linear interpolation in the shape space, to model the dynamics of cell differentiation using only static images. Without any prior knowledge, the method produces a realistic shape evolution process.

**Author Summary**

Cellular differentiation is an important process that is challenging to study due to the number of organizational changes it includes and the different time scales over which it occurs. Fluorescent microscopy is widely used to study cell dynamics and differentiation, but photobleaching and phototoxicity often make it infeasible to continuously observe a single cell undergoing differentiation for several days. In this work, we described a method to model aspects of the dynamics of PC12 cell differentiation without continuous imaging. We constructed accurate representations of cell and nuclear shapes and quantified the relationships between shapes and mitochondrial distributions. We used these to construct a generative model and combined it with a matching process to infer likely sequences of the changes in single cells undergoing differentiation.

## Introduction

Cellular differentiation is a highly complex process that is incompletely understood. While fluorescence microscopy provides a widely-used tool for investigating the organization of cell components, given the number and complexity of the resulting images it is clear that there exists a need for automated methods for their analysis [1]. Tools are needed not just for describing these images, but also for creating models of cell organization that incorporate information from many cells [2].

Due to the intimate relationship between neuron morphology and function, particular attention has been paid to how to model and represent cell shapes. Tools have been described for tracking neurites [3] and modeling neuronal structure [4-6] using segmented electron or fluorescence microscope images. While some methods are primarily concerned with representing neuron shape via summary statistics such as shape and skeleton features [6], the software L-NEURON and ARBORVITAE [4] use distributions over semi-parametric tree representations to construct generative models of neuron morphology capable of synthesizing cell shapes. The NETMORPH software [5] likewise uses a generative modeling framework, but is additionally capable of constructing large networks of interconnected cells. Modeling of the dynamics of cell shape and organization during processes such as differentiation has received less attention. Using continuous imaging to construct models of single cells throughout a differentiation process is difficult due to compounded effects of both phototoxicity and photobleaching and the difficulty of tracking individual cells from sparse time points in long time series. Furthermore, modeling correspondences between individual neurites at different timepoints in the parametric models previously proposed is not straightforward, and those models represent cell shapes as tree structures without considering neurite or cell body thickness. Although the neuron generation procedures proposed could be interpreted as growth models, they are described as biologically inspired rather than learned directly from time series.

Mitochondria have been shown to have a role in cell differentiation fate [7], but their spatial distributions are difficult to represent due to the fact that they form complex dynamic networks. Furthermore, there has been little work on describing the relationship between cell morphology and mitochondrial distribution.

Of the many model systems for cell differentiation, rat pheochromocytoma cell line PC12 is particularly useful for studying neuronal differentiation and survival [8-10]. After stimulation with Nerve Growth Factor (NGF), PC12 cells differentiate into sympathetic neuron-like cells, a process which is morphologically marked by neurite outgrowth over a time course of up to six days [11-14]. To address the goal of building continuous models of cell shape and mitochondrial distribution during differentiation, we collected images of PC12 cells at various times after treatment with NGF. From these we constructed a joint cell and nuclear shape model based on spherical harmonic descriptors [15] and a probabilistic model of mitochondrial localization [16] and combined them into a generative model of shape and mitochondrial distribution over all time points. We then developed a novel approach for combining these models to predict likely sequences of changes that single cells undergo through the differentiation process despite the fact that movies of single cells were not available.

## Results

### Cell Component Representation

As described in the Methods, we collected 3D images of mitochondrial staining of PC12 cells at various times after treatment with NGF. This was done in two large experiments, one consisting of images every 12 h up to 48 h, and one every 24 h up to 96 h; the experimental setup is illustrated in S1 Fig. We decomposed the image of each cell into three components: a cell shape, a nuclear shape and a mitochondrial spatial distribution. Fig 1 shows this procedure on a typical cell image.

**Fig 1.**
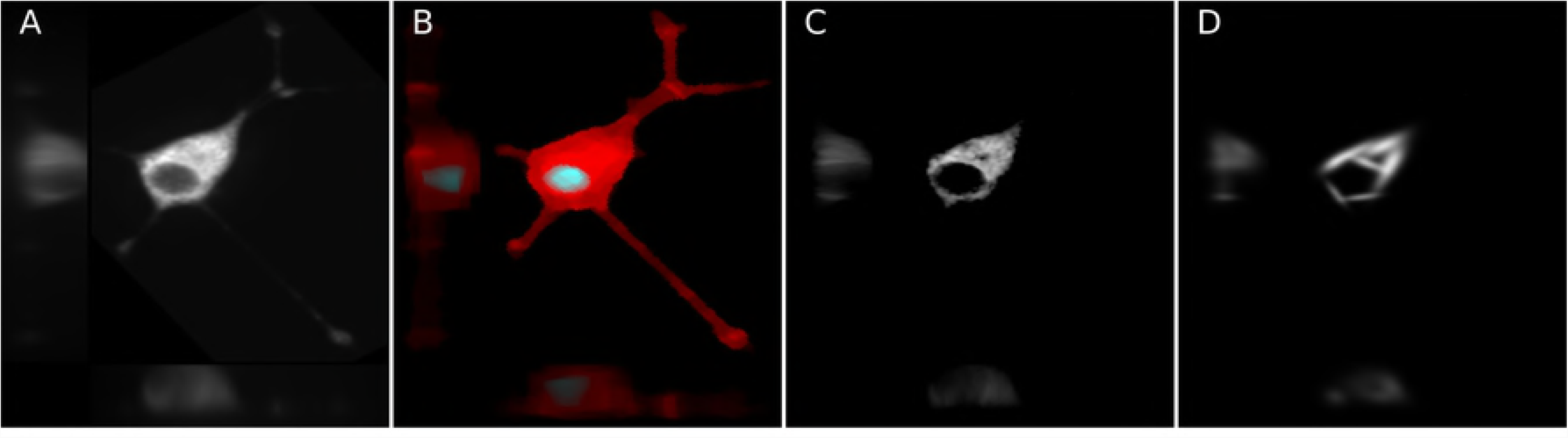
Per-cell shape and mitochondria localization modeling procedure. The image was segmented into cell and nuclear shapes and these were aligned using the SPHARM-RPDM method. An aligned original image (A) and the segmented cell (red) and nuclear (gray) shapes (B) are shown; these are used to create a shape space model. The individual mitochondrial objects from the original image are found using a Gaussian mixture model (C), and their positions are modeled as a probability density function (D).

### Models of Cell and Nuclear Shape

Cell and nuclear shapes were first converted into spherical harmonic descriptors using Robust SPHARM-PDM (SPHARM-RPDM) [15] with shape alignment and scale normalization as described in the Methods. Out of 997 cells in the original dataset, 8 cells that were not well represented by the descriptors were removed from the analysis. Dimension reduction was done on the descriptors using principal components analysis (PCA) to generate a specified number of latent features. We found that models constructed with 300 dimensions were able to capture the cell shapes of individual cells with high accuracy, as shown in S1 Table. Some examples of reconstructed shapes from the models with the corresponding original shapes are shown in S2 Fig. The models were constructed with two different methods of shape alignment, using the first order ellipse as done previously [15] or using the major axis (see Methods). The reconstructions errors were similar, but since they were slightly better for the major axis alignment approach, all subsequent analysis was done using that method.

To provide a loose illustration of the major trends in shape as a function of differentiation, low dimensional shape spaces constructed from the latent features are shown in Fig 2 and S3 Fig with or without the scale factors that were removed during the initial normalization. Cells from different time points overlap in shape fairly extensively, but there is a trend towards an increase in size and in the first shape component (PC1, which corresponds approximately to elongation); this is consistent with previous observations that PC12 cells start from a roughly spherical morphology and gradually flatten and spread out with more and longer neurites (that is, with more complex cell shapes) after NGF treatment. It is important to note that these two-dimensional representations do not allow full visualization of the cell and nuclear shape variation. The first principal component captures 33.6% of that variation and the second captures 7.9%, leaving 58.5% unvisualized in these two-dimensional maps. However, all operations using the models described below were done in the high dimensional shape space.

**Fig 2.**
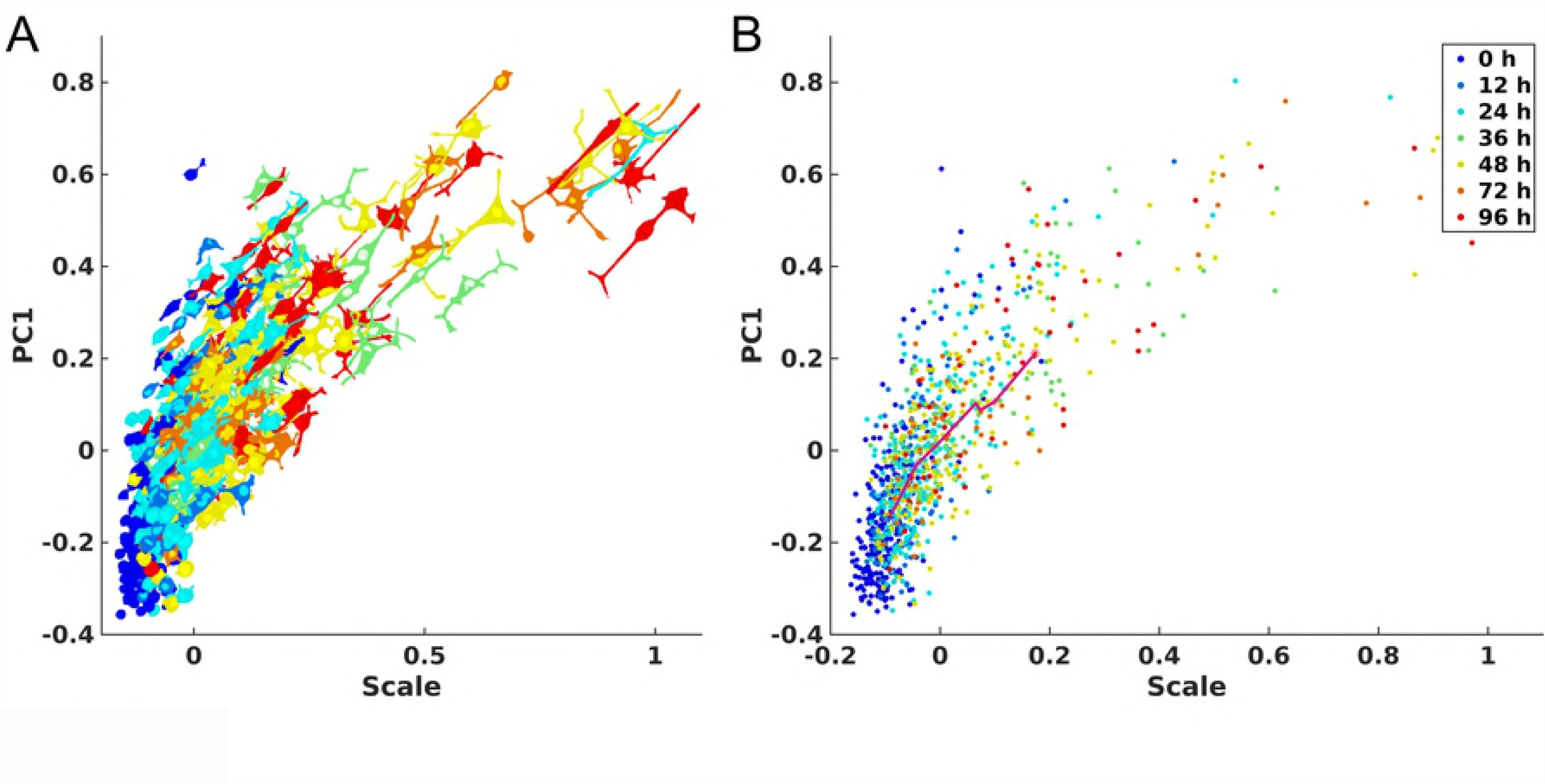
Shape space for the joint model of cell and nuclear shapes constructed from all cells for all the time points in the two experiments. Here the cell size and first principal component are shown. Panel A shows the space with images projected in the XY-plane in the corresponding locations. Panel B shows a scatter plot with points for each cell shape; the line links the centroids of the points for adjacent time point to indicate the trend as differentiation proceeds. In both panels, blue indicates untreated cells and warmer colors indicate later time points.

### Relationship between mitochondrial localization and cell and nuclear shape

For each cell in the collection, the distribution of mitochondrial localization was described as the probability of a mitochondrial object occurring at a position inside of the cell according to a standardized coordinate system relative to the cell and nuclear membranes. We used the CellOrganizer implementation of the previously described method [16] in which each object is represented by its relative distance from the nucleus and the azimuth and angle from the major axis and the positions of all objects are fit using a logistic model (see Methods). The mitochondrial distribution for each cell is thus represented by the 6 parameters of the model. Given these parameters, we asked how the relationship between the mitochondrial location pattern and the cell shape changes as a function of differentiation.

To evaluate this relationship, we used multi-response regression to predict the mitochondria localization model given the cell and nuclear shapes, as described in Methods. We used nested leave-one-out cross validation to first determine the optimal regularization parameters λ_1_, *λ*_2_ and *λ*_3_ and the corresponding model parameters 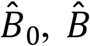The parameters of the held-out cell were predicted, and the error between the predicted and measured mitochondrial parameters was recorded (this error serves as an inverse measure of the extent to which cell shape and mitochondrial localization pattern are related). Boxplots illustrating the distribution of errors at each time point and experiment with or without scale factor are shown in S4 Fig and Fig 3. There is a distinct trend towards a decrease in the error of predicting the mitochondrial localization pattern as a function of time after treatment. We compared the errors between treated time points with the initial time point without treatment (0h) via the t-test and corrected for multiple tests using Bonferroni-Holm correction [17]. An asterisk indicates a significant difference in the ability to predict the mitochondrial location pattern from the cell and nuclear shape between this time point and 0h. As can be seen in Fig 3 for predictions with only shape models, the prediction errors decreased significantly over time, compared to those in the initial untreated condition. Also, the decrease is most dramatic in the beginning (12h for 48-h experiment, 24h for 96-h experiment). For the model with scale included, the patterns of prediction errors are similar, as shown in S4 Fig. The similarity between results for models with or without scale suggests shape variation rather than cell size is dominant in the prediction of mitochondria pattern.

**Fig 3.**
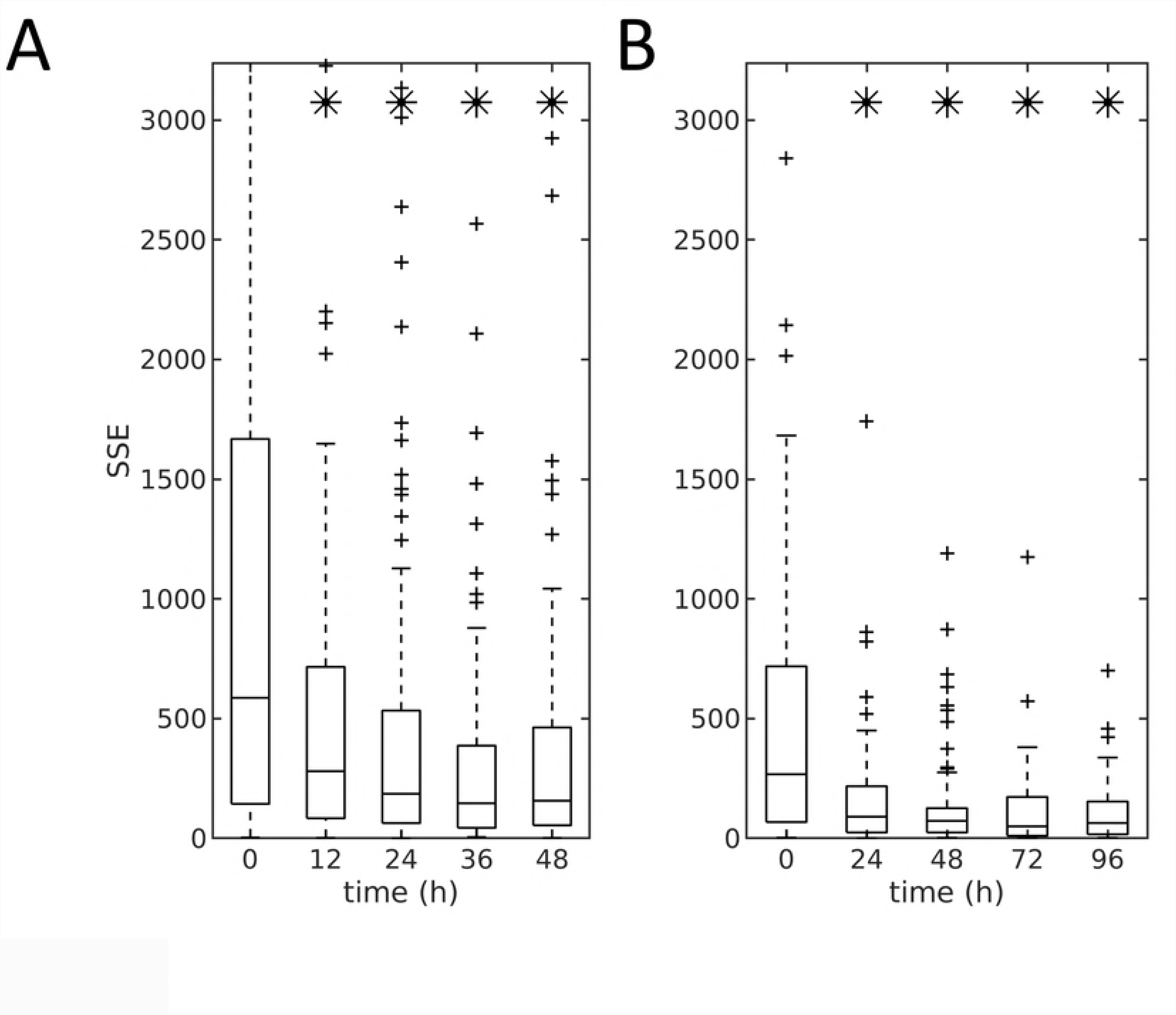
Prediction error of mitochondrial localization parameters as a function of time for the model between shapes (without size) and mitochondria patterns. Panels A and B show the results for the 48-hour and 96-hour dosing experiments, respectively. At each time point (x-axis) the central box mark indicates population median, and the lower and upper bounds of the box indicate 25^th^ and 75^th^ percentiles. Whisker bounds cover approximately 95% of the data, with outliers shown in small crosses. An asterisk indicates that the error for that time point is statistically different from the error at the 0h time point.

For the decrease of the prediction errors across time, one potential explanation could be that the variation in the mitochondrial distribution from cell to cell decreases with treatment time (and thus predicting a close mitochondrial distribution is made easier). To test this, we determined whether the errors for a mitochondrial distribution predicted from a cell’s shape space position were significantly smaller than those resulting from random choice of a cell from all cells in a given experiment. The models were all significant at *α* < 0.05 after Bonferroni-Holm correction as shown in S2 Table. These results indicate that a significant relationship exists between mitochondrial localization and cell shape and that the relationship becomes stronger as a function of time.

Fig 4 shows the distributions of the parameters of the mitochondria model for each time point for the 48h and 96h experiments. B_1_ and B_2_ (parameters weighting the distance from the nucleus) show a strong relationship to time after treatment; they also show a high degree of correlation (Fig 4B), becoming more constrained as a function of time after treatment. To illustrate variation in mitochondrial patterns across time, S5 Fig shows example cell shapes, segmented mitochondria patterns, and modeled and predicted spatial probability density models, for average cell shapes every 24h for the 96h dataset.

**Fig 4.**
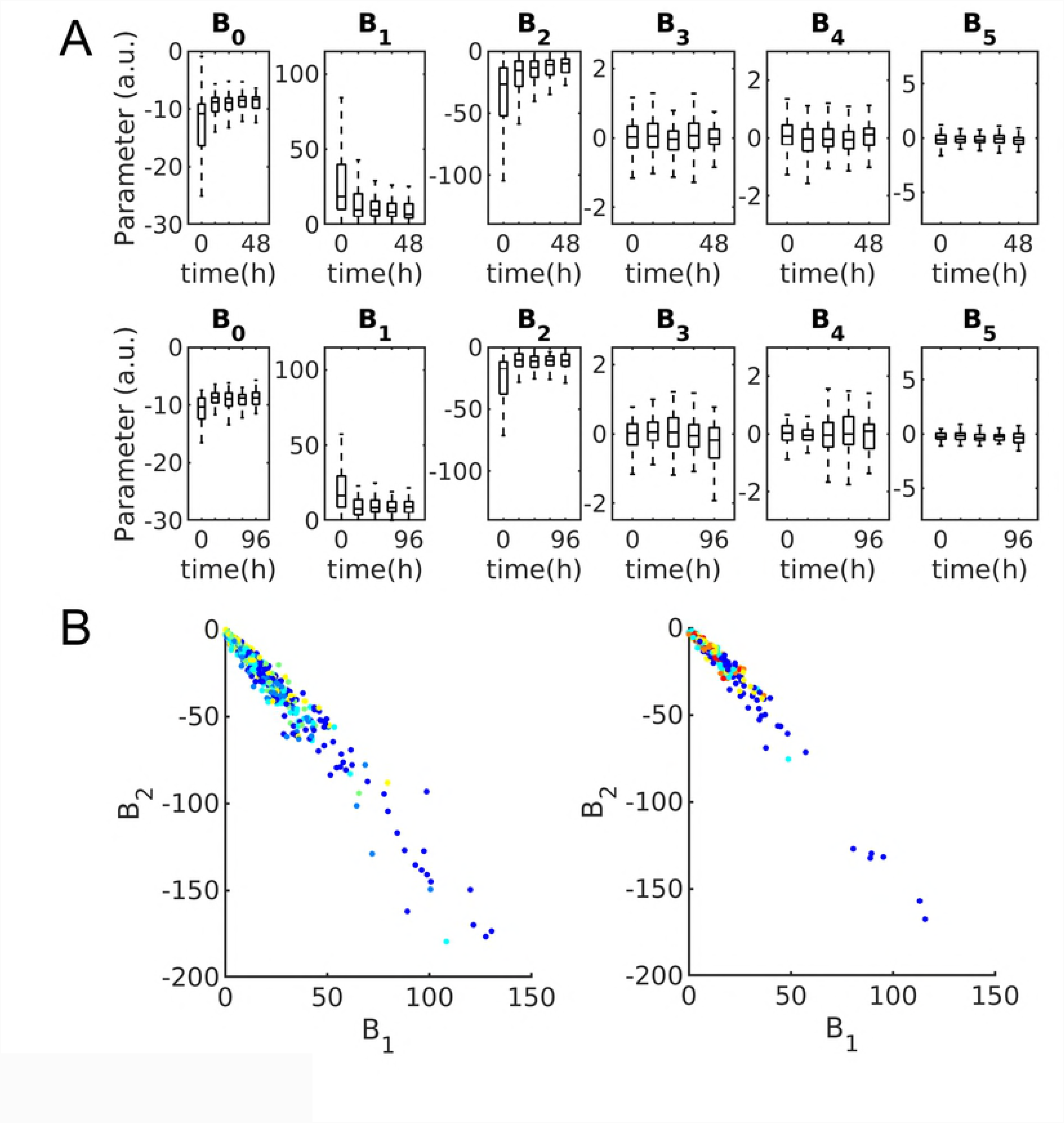
Mitochondria distribution parameters. A) Boxplots showing distribution of parameters for 48-hour (top) and 96-hour (bottom) experiments. Box centers indicate population median with bounds indicating 25^th^ and 75^th^ percentiles respectively. Whiskers indicate 99% coverage of data. B) Mitochondria distribution parameters corresponding most with time plotted against each other and colored by time for 48-hour (left) and 96-hour (right) experiments. Blue indicates untreated cells and warmer colors indicate later time points at either 12-hour or 24-hour intervals.

### Modeling kinetics of differentiating cells

We next sought to construct a model of shape dynamics, such that we could generate movies of synthetic shapes for cells as they differentiate. Since we do not have images of the same cell at different time points, we cannot directly learn a dynamic model using the approach we have previously described [18]. Here we therefore propose an alternative model for shape dynamics. The basic idea is to assume that the populations of cells at each time point are large enough that we can consider that for each cell in our collection for a given time point that there is a cell in the collection for the next time point that is reasonably similar to what the first cell would have looked like at the later time. We find the matches between cells at adjacent time points that give the lowest total difference in shape between them (by weighted maximum bipartite matching, as described in the Methods). This gives us a “trajectory” in time and in shape space for each cell at the 0h time point (without NGF treatment). Using shape evolution (synthesis) [15] we can construct intermediate shapes within each trajectory by interpolating along a linear path in the shape space between each pair of shapes in adjacent time points.

The expected shape differentiation models are shown in S6 Fig and Fig 5 for the 48-hour and 96-hour experiments respectively, with finer and smoother synthesis of corresponding trajectories shown in S1-S8 Videos. In both figures, each row shows the evolution of cell and nuclear shape for a given cell from 0h to 48/96h. The four cells are chosen based on quantiles of total distances between the matched cells of adjacent time points across all time points. From the figures, we can see that the shape evolution method appears reasonable in terms of the reconstructions of cell shapes for either observed or unobserved cells (interpolated time points), and captures the expected trend from round cells to complicated shapes with long neurites (for most trajectories). Also, the total distances in the shape space for the trajectories reflect the overall shape variance across time, e.g. the final shapes generally become more and more complicated as the quantile increases. Moreover, the sensitivity to NGF treatment is clearly heterogeneous among PC12 cells, as some of the matched cells do not differentiate after treatment (the presence of these cells in the late time points of course indicates this as well). In S7 and S8 Fig, the expected directions for the transitions of cell shapes for different size time steps are shown. The figures confirm the last observation, as different positions in the shape space are predicted to move towards quite heterogenous directions at the next time point (especially during the early stages). This finding of heterogeneity agrees with previous experimental studies [9, 19].

**Fig 5.**
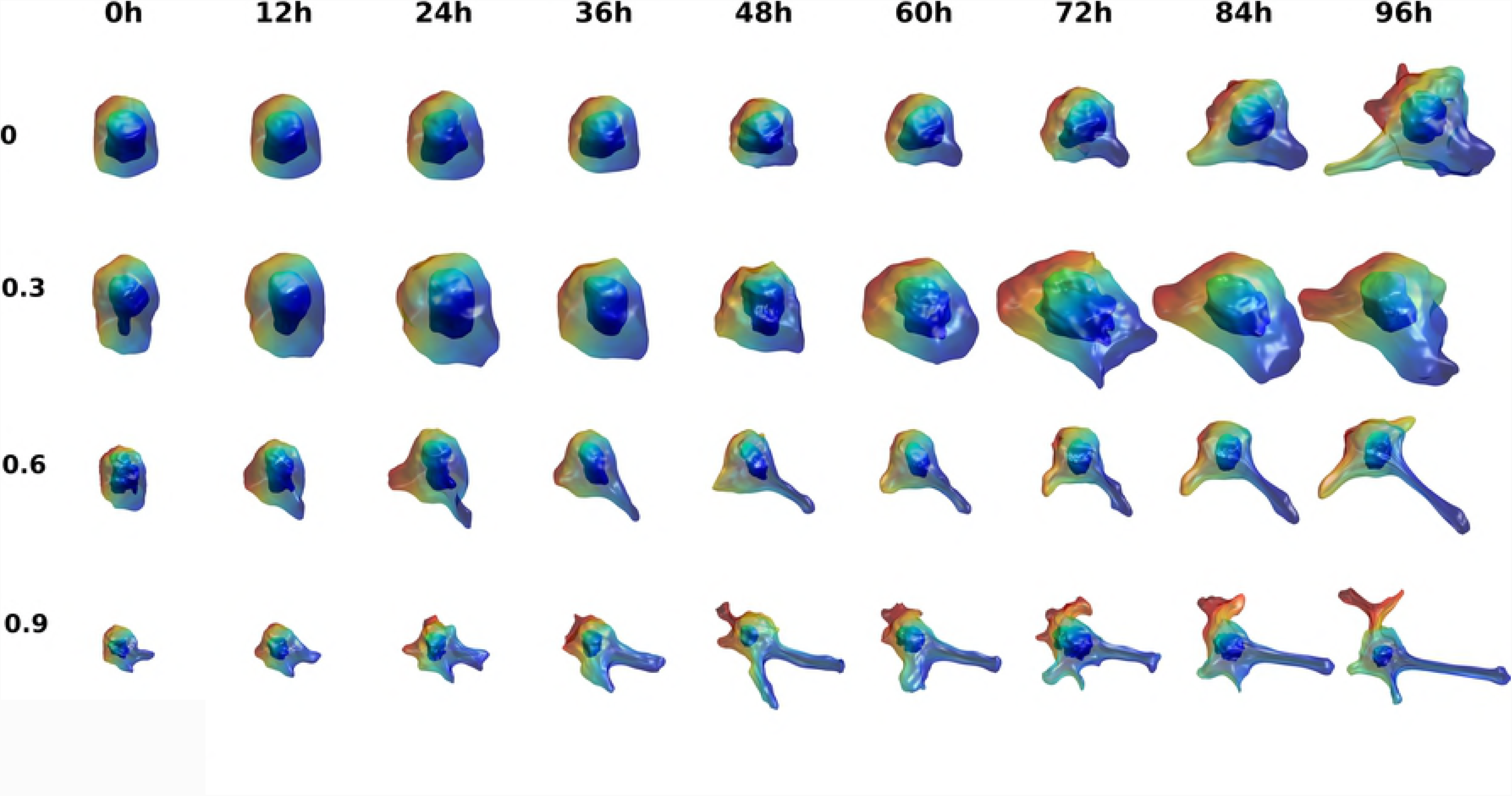
Illustration of cell and nuclear shape differentiation for 12-hour time steps given cell shape 96-hour dosing experiment. Four trajectories were chosen based on the quantiles of total distances between each matched cell pairs in the trajectory. The quantiles are shown in the y-axis. The time points are shown in the titles. Time points 12h, 36h, 60h and 84h are interpolated using the cells in the previous and following time points.

### Generative model of the relationship between cell shape and mitochondrial location pattern

In addition to simulating dynamics for cell shape, we can also model the dynamics of changes in mitochondrial distribution. Using the regression model between cell/nuclear shapes and mitochondrial patterns, we inferred parameters of the mitochondria model for each of the interpolated cell and nuclear shapes along our estimated cell trajectories. This allows a probability density distribution for mitochondria to be synthesized using the inferred parameters and the volume images of cell and nuclear shape (using the image synthesis function in CellOrganizer). Fig 6 shows the mitochondrial probability densities for the same four cells as in Fig 5. The complete sets of frames of mitochondrial patterns for these 4 cells are shown in S9-S12 Videos. Figs 5 and 6 illustrate that the method can generate very realistic intermediate cell shapes, even though there is no prior knowledge or constraints on the unobserved shapes. More importantly, the shape evolution process is also realistic, in that it represents the neurite growth process in a reasonable manner, even though no prior knowledge of how the neuron develops is provided.

**Fig 6.**
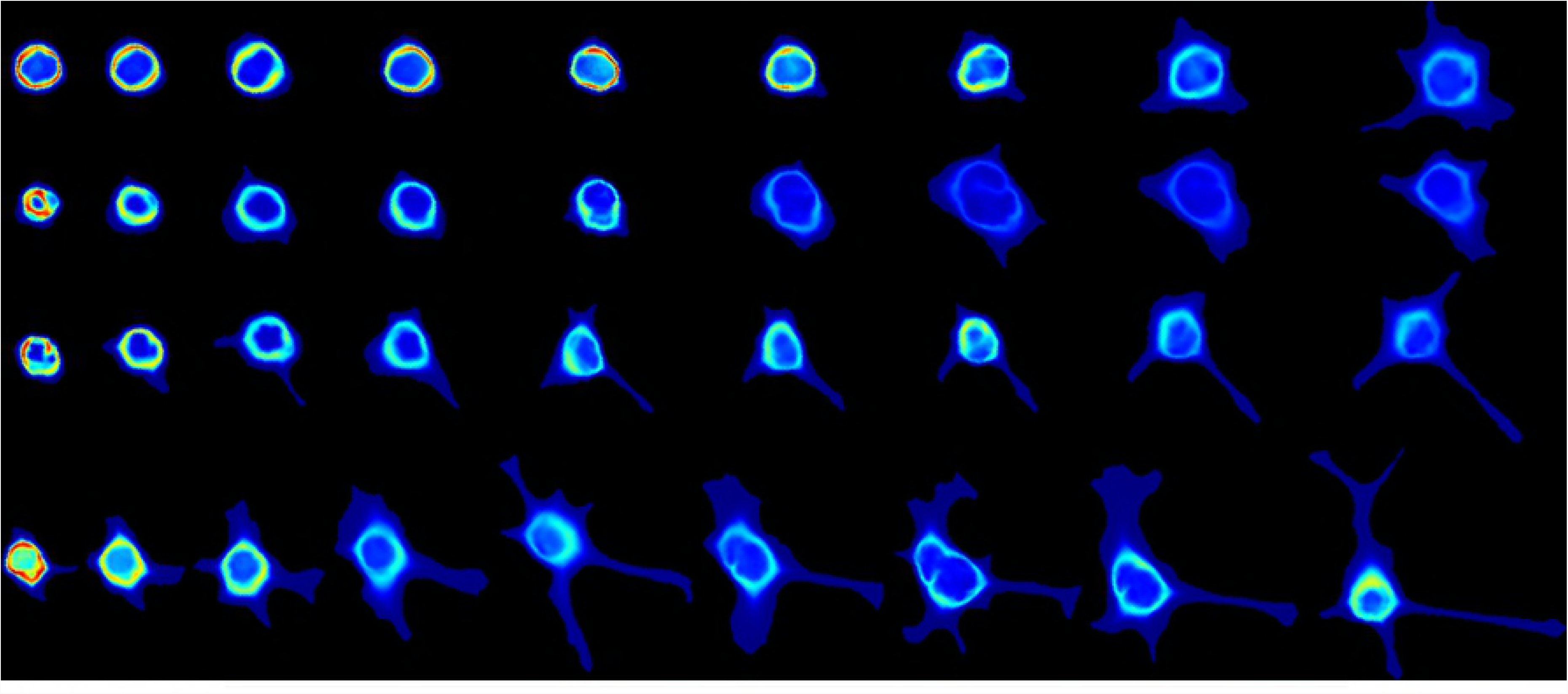
Illustration of mitochondria localization patterns in the differentiation for 12-hour time steps in the 96-h experiment. Each row shows 2D mean value projections of the 3D mitochondrial probability density maps in a trajectory. The trajectories, as well as time points are the same as shown in Fig 5. Each column represents a time point from 0h to 96 h with time step of 12h from left to the right. The maps are shown with a hot-cold color map (blue indicates low probability of observing a mitochondrion at that location).

## Discussion

One of the objectives of systems biology is to understand the relationships between cell compartments in a manner such that cell fate and the organization of unobserved components may be predicted. An additional objective is that these models not rely on human interpretation, such as hypothesized mechanisms for biochemical processes, but rather be learned directly from experimental measurements.

With these considerations in mind, we developed a tool to model the relationship between cell morphology and organelle organization, and demonstrated that this relationship varies during differentiation of PC12 cells. We found that there is a decrease in variation of mitochondrial localization with respect to time after differentiation.

Given population snapshots we constructed a model to describe shape evolution in response to NGF treatment; the model is capable of producing movies by statistically sampled differentiating cell shapes. Here we make very simple assumptions for shape dynamics in terms of both cell/nuclear shapes and mitochondria localization by interpolating across linear paths in the shape space for the cell and nuclear shape models and then estimating parameters for the localization model from those shapes. The synthetic movies appear reasonable in terms of shape dynamics even without using any prior knowledge of how PC12 cells actually differentiate. Of course the model of mitochondrial localization is quite simplistic, only considering the spatial probability distribution, rather than trying to predict individual mitochondrial shapes, sizes and intensities. Thus, one potential future direction is to apply other generative methods for organelle or protein dynamics, for example, the 3D equivalent to optimal transport models [20].

The method we have described is capable of constructing models in a range of time-varying cell-component localization applications, including but not limited to changes associated with division and cell migration. Recent findings have shown population heterogeneity is inherent in the PC12 signaling networks [21]. The relationship between proliferation and differentiation is sharply defined by mutually exclusive pAKT and pERK concentrations [22]. This suggests an influence of stochastic effects on the cell fate decision, and that on the single cell level cells are either proliferating or differentiating. As a consequence, homogeneity of the population can be reduced by optimal growth conditions, but never completely abrogated [23, 24]. The image-derived modeling technique described here is able to model single-cell decisions and is therefore a small step in the development of tools to automatically discover relationships between cells and their components, as well as provide compact representations of these relationships learned directly from images.

## Materials and Methods

### Cell Culture and Experimental conditions

PC12 cells (between 6 and 10 passages) were obtained from ATCC (American Type Culture Collection, UK) and were cultured in RPMi medium containing 10% horse serum (HS), 5% fetal calf serum (FCS) 1% L-Glutamine and Penicillin/Streptomycin at 37°C in 5% CO_2_. Cells were plated on collagen coated 35mm glass-bottom ibiTreat dishes and were allowed to adhere for 24 hours. Two types of experiments were performed. Cells were either treated with 50ng/ml rat Nerve Growth Factor (NG; Promega, Madison, WI, USA) at 0, 12, 24, 36 and 48h prior to imaging at the same time, or were treated at the same time and imaged at 24 h increments up to 96 h after treatment. An hour prior to imaging, cells were stained at 37° C with a 0.5uM solution of Mito Red (Sigma-Aldrich, Munich, Germany) for 5 minutes, rinsed with PBS and placed in 1ml of growth media without Phenol red.

### Microscopy

Cells were imaged on an Axio Observer.Z1 (Carl Zeiss Microscopy, Jena, Germany) microscope equipped with a spinning disk (CSU-22, Yokogawa, Japan) with an EX-Plan-Neofluar 40x/1.30 Oil objective. The sample voxel size was 0.161 um x 0.161 um x 0.340 um and 59 slices were taken with a 150ms exposure time at 12-bit pixel depth. Imaged cells were manually selected to not be in contact with other cells. Due to the sensitivity of cells to phototoxicity, approximately 10 fields were imaged per plate. Between 172 and 98 cells were imaged at each time point for the 48h experiment and between 46 and 89 cells per time point in the 96h experiment.

### Cell Shape Segmentation

Each slice of an image was convolved by a 2D Hessian filter of 3 standard deviations and Eigen-edges were extracted [25]. Dilate and erosion operations were performed on each slice with a disk structuring element of 14 and 24 pixels respectively. The final shape was regularized by convolving with a Gaussian of 7-pixel standard deviations and retaining all pixels with a value greater than 0.5. The result was a “shell” of the cell shape, and thus a fill operation was performed on each remaining region.

### Nuclear Shape Segmentation

Given a masked cell shape, the intensity image was thresholded via Ridler-Calvard [26] thresholding. The nucleus is treated as “not signal” in the thresholded image within the region of its convex hull. Because this may result in spurious objects, a distance transform was performed, segmented with an active contour, and the largest object returned as the final nuclear shape. An example of this pipeline is shown in S9 Fig.

### Cell and Nuclear Shape model with spherical harmonic framework

#### Shape alignment and modeling

Joint models for cell and nuclear shapes were constructed using the spherical harmonic framework as described [15]. To make different shapes comparable, this framework aligns shapes using the first-order ellipse before creating the model. As an alternative we also did alignment using the major axis. For this, the primary direction was obtained by projection of surface points to the XY-plane, followed by PCA to find the major axis. The cell shape was then aligned to this axis. After that, if the skewness along the x-axis was negative, the shape was flipped in the XY-plane.

After alignment of the object in the original space, it is also necessary to align the parameterization so that the final descriptors of different cells are comparable. The basic idea is to find some landmarks from the parameterization in specific directions in the original space, such as poles, points in the equator, and then rotate the parameterization. Here as a first step, a pair of vertices whose direction is mostly close to x-axis is picked as poles in the spherical parameterization. To do so, first, vertices in the object are paired with each other such that the projection to the unit sphere of one point is closest to the antipodal point of that of other point (that is, the two points are (approximately) diametrically opposite to each other after mapping). Second, pairs of matched points whose vector directions are almost within XY-plane are selected as candidate north and south poles with first 1% in terms of smallest angles to XY-plane. Then from these pairs, a subset of pairs is chosen if the absolute values of the x-axis in the direction vectors are greater than a threshold (0.9999 in our implementation), in order to pick out those pairs whose directions are almost closest to the x-axis. After that, a pair with longest distance in this subset is established as south and north poles.

After finding the poles, the landmarks in the equator (similar as points with 0^°^ and 90^°^ longitudes in the equator) need to be picked out. First, based on the poles, the pairs of points with θ angles mostly close to the zero are chosen. Among these points, the pair with minimum differences in the z-axis is chosen as landmarks for equator. The rotation matrix is defined as the rotation from the projected coordinate of the equator landmark (in the parameter space) with larger y-coordinate to the coordinate of (0, 1, 0). After rotation, the spherical parameterization will be flipped along x-axis, if the point in the object space with coordinate (1, 0, 0) in the parameter space has a smaller z-coordinate than that of the centroid of all surface point.

#### Shape reconstruction

Shape reconstruction from a SPHARM-RPDM model was done as described previously [15]. The accuracy of shape reconstruction was measured using Hausdorff distance, which is defined as

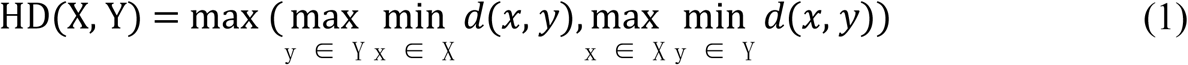

where X and Y are two sets of points, and *d*(*x, y*) is a metric of distance between two points (Euclidean distance in our case). The 3D volume images of shapes were converted to surface meshes, and vertices in the meshes for the original and reconstructed surfaces were used to calculate the Hausdorff distance. An additional error metric, peak signal-to-noise ratios (PSNR) between the original and reconstructed shape, was also included to evaluate the reconstruction quality. PSNR is calculated based on the Hausdorff distance with the following form:

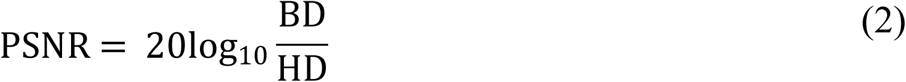

where BD is the diagonal of the minimum bounding box of the cell and HD is the Hausdorff distance. For the joint model, the joint reconstruction error was defined as the average of those for the two components (cell and nuclear shapes).

#### Mitochondrial Localization Model

Mitochondrial localization models were learned as described previously [16]. Briefly, the mitochondrial image after masking to the cell boundary was preprocessed by removing intensity below the Ridler-Calvard threshold. A spherical Gaussian mixture model was fit using seeds at each intensity local maxima after convolving the image with a Gaussian filter of one voxel standard deviation. The position of each voxel in the cell was parameterized according to its ratio of distance to the nuclear surface over the nuclear distance plus the distance to the cell surface, *s*, and the inclination and azimuth angles, *θ* and *ϕ* respectively, from the nuclear center, and logistic function was fit to the probability that each pixel contains an object centroid,

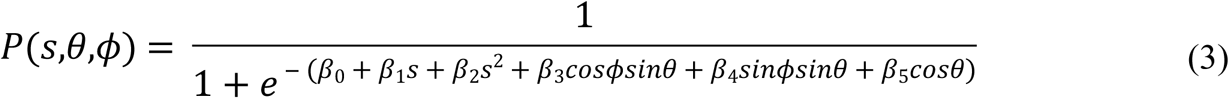

The spatial probability distribution for each cell was parameterized by the 6-element vector *β.*

### Regression model between shape and mitochondrial distribution

We used a multi-response regression to model and predict the mitochondrial localization model given the cell and nuclear shapes:

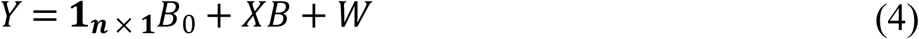

Where *X* ∈ *R*^*n*×*s*^ is a matrix of joint shape-space positions of dimension *s*, with each row corresponding to a cell and nuclear shape, and each column a dimension of the shape space (in this case 300 dimensions without scale, 301 dimensions including scale factors as an additional feature). *Y* ∈ *R*^*n*×*p*^ is a matrix of mitochondrial localization models, with each row being a model corresponding to the cell at the same row in *X*. 1_*n* × 1_ is the n-dimensional column vector with all elements as 1. *B*_0_ and *B* are model parameters, where *B*_0_ ∈ *R*^1 × *p*^ is the parameter for the intercept and *B* ∈ *R*^*s* × *p*^ is the regression matrix describing the relationship between the shape space and mitochondria localization models. *W* ∈ *R*^*n* × *p*^ is a matrix of random noise following multivariate Gaussian distribution with zero-mean (the residual variation in the localization parameters not explained by model parameters). Here we combined elastic net regression [27] with a group-penalized estimator [28] of model parameters defined as

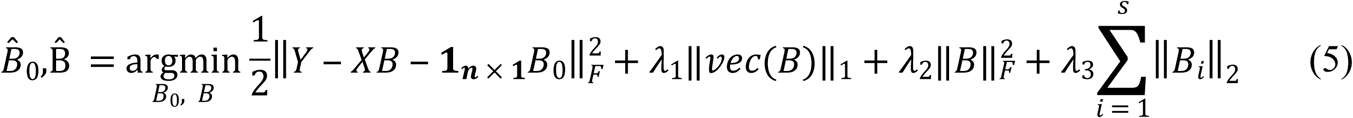

where *vec*(.) is the operator of reshaping a matrix into a column vector, ∥ . ∥_1_ represents the *l*_1_ norm of a vector, ∥ . ∥_*F*_ stands for the Frobenius norm of a matrix, and ∥*B*_*i*_ ∥_2_ indicates the *l*_2_ norm of i-th row of *B*. The regularization parameters *λ*_1_, *λ*_2_ and *λ*_3_ function as penalization terms on *B* to control the structure of *B*, as well as to avoid overfitting of the model. These regularization parameters were chosen by sweeping over combined sets of possible candidate values, and selecting the set that results in the lowest mean-squared prediction error of *Y* via 10-fold cross validation. In the cross validation, we allowed λ_2_ and/or λ_3_ to be zero, which means that the model may degenerate into lasso regression [29] (if both λ_2_ and λ_3_ are zeros), elastic net regression [27] (if λ_3_ is zero), or sparse group lasso regression [30] (if λ_2_ is zero). The reason for the possibility of degeneration in the model is to allow more flexible control of the model in response to different situations in the datasets. We implemented the regression model with the ADMM (alternating direction method of multipliers) framework [31].

### Modeling kinetics of differentiating cells

Given cell populations at sequential time points, we sought to find plausible and most similar cell shapes at the subsequent time point; such a shape-pair can be treated as a “trace” in time series models. This essentially becomes a matching problem, where we want to find a matching of each cell in one time point to one cell in the next time point that minimizes the total shape space distance between pairs of matched cell shapes. More formally, given the shape space positions of equal numbers of cells at subsequent time points, 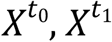, we can construct a matrix of cell shape distances between cells at subsequent time points,

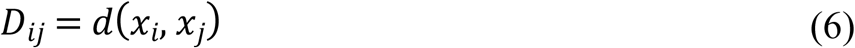

Where 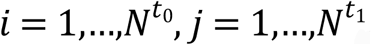, and 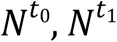 are number of cells for *t*_0_ and *t*_1_, respectively. We want to minimize the function

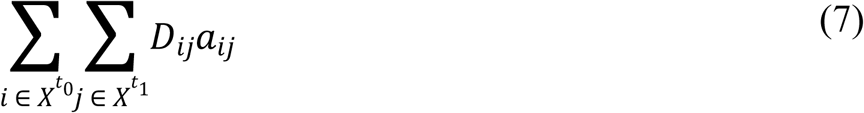

where *a*_*ij*_ is binary matrix of assignments taking a value of 1 if there is an assignment, and 0 otherwise, subject to the constraints 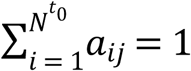 and 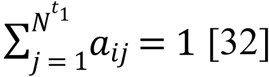. This problem was solved through the Hungarian algorithm [33].

## Acknowledgments

This work was supported in part by U.S. National Institutes of Health grants GM090033, GM103712 and EB009403, by the Excellence Initiative of the German Federal and State Governments through the Freiburg Institute for Advanced Studies, and by a Research Award to RFM from the Alexander von Humboldt Foundation.

## Availability

The CellOrganizer software used here for modeling these relationships is available at http://cellorganizer.org. The source code for performing all analyses in this paper, as well as analysis results, is available for reviewers at http://murphylab.cbd.cmu.edu/software/2019_PC12. This link is currently hidden (and not tracked) and will be made public upon manuscript acceptance by linking to it from http://murphylab.cbd.cmu.edu/software. The original images are available at https://datadryad.org/review?doi=doi:10.5061/dryad.hc8037v

## Supporting information

**S1 Fig. Treatment and imaging protocol for PC12 cells.** Two experiments were performed. 24 hours after plating, cells were treated with NGF at 12 or 24-hour intervals for 48 or 96 hours, each resulting in 4 time points after NGF treatment and an untreated group.

**S2 Fig. Illustration of shape reconstructions with the SPHARM-RPDM model.** Panels A and B show reconstructions from a model of only cell shape and both cell and nuclear shapes, respectively. The original shapes, reconstruction with the model using alignment by the major-axis method, and reconstruction with the model using alignment by FOE method are shown. For both cases, the reconstructions are generated using models with 300-dimensional representations. For each panel, the cells in the columns were picked based on the quantiles of Hausdorff distances of the major-axis method (with the quantile shown at the top of the column); thus the cells on the left were easier to reconstruct and the reconstructions of cells on the right reveal some minor differences. The HD and peak signal-to-noise ratio (PSNR) are listed under each reconstruction.

**S3 Fig. Shape space for the joint model of cell and nuclear shapes constructed from all cells as all time points in the two experiments.** Here the first two principal components are shown. Panel A shows the space with images projected in the XY-plane in the corresponding locations. Panel B shows a scatter plot with points for each cell shape; the line links the centroids of the points for adjacent time point to indicate the trend as differentiation proceeds. In both panels, blue indicates untreated cells, where warmer colors indicate later time points, as shown in the legend of panel B.

**S4 Fig. Prediction error of mitochondrial localization parameters as a function of time for the model between shapes including cell size and mitochondria patterns.** Panels A and B show the results for the 48-hour and 96-hour dosing experiments respectively. At each time point (x-axis) the central box mark indicates population median, and the lower and upper bounds of the box indicate 25^th^ and 75^th^ percentiles. Whisker bounds cover approximately 95% of the data, with outliers shown in small crosses. An asterisk indicates that the errors for that time point are statistically different from the errors at the 0h time point.

**S5 Fig. Example mitochondrial distribution patterns for most-average cell shapes at 24-hour intervals for the 96-hour experiment.** Each column shows cell shape, segmented mitochondrial pattern, modeled mitochondria spatial probability distribution, predicted mitochondria spatial probability distribution, and the probability distributions registered to a “canonical” cell shape of two nested spheres with the volume of the average nucleus and cell sizes respectively.

**S6 Fig. Illustration of synthetic cell/nuclear shape differentiation for 6-hour time steps given cell shape 48-hour dosing experiment.** Four trajectories are chosen based on the quantiles of total distances between each matched cell pairs in the trajectory. The quantiles are shown in the y-axis. The time points are shown as in the titles. Time points 6h, 18h, 30h and 42h are interpolated using the cells in the previous and following time points.

**S7 Fig. Expected shape differentiation estimated from trajectories in the 48h experiment for 12-hour time steps.** The expected shape change is estimated according to the matching between each pair of time points. Color indicates relative probability density of a cell shape at that time point, and the vector indicates the direction of the shape change. White dots indicate positions of observed shapes. The remaining shape-space dimensions have been marginalized out. Vector arrows have been scaled for visualization purposes.

**S8 Fig. Expected shape differentiation estimated from trajectories in the 48h and 96h experiments for 24-hour time steps.** The first row shows the expected transitions from 0h to 24h and 24h to 48h in the 48h experiment as comparisons to those in 96h experiment. The expected shape change is estimated according to the matching between each pair of time points. Color indicates relative probability density of a cell shape at that time point, and the vector indicates the direction of shape change. White dots indicate positions of observed shapes. The remaining shape-space dimensions have been marginalized out. Vector arrows have been scaled for visualization purposes.

**S9 Fig. Nucleus segmentation procedure.** The original image (A) was segmented (B), and a convex hull was formed (C). The candidate nuclear region is the “not signal” from the segmented image within the convex hull (D). The result is distance transformed (E) and segmented via active contour methods (F).

**S1 Table. Reconstruction errors for cell shape and joint models with FOE and Major-axis alignment.** To calculate reconstruction errors, the dataset was randomly split into training (842) and testing (147) sets, with the model built with training set and the reconstruction error calculated over the testing set. The cell shape model and joint model were built with either FOE alignment as described [15] or major-axis alignment as described in Methods. The number of latent dimensions for both models is 300. The means of HD and PSNR (in parenthesis) as defined in Methods among cells in the testing set are shown here. Lower HD and higher PSNR indicate better reconstructions. For joint modeling, the errors for cell shape, nuclear shape and average of the two components (joint error) are all shown in the table for reference.

**S2 Table. P-values of t-tests of prediction errors for mitochondrial distributions from shape models.** For each experiment and each condition (with or without including of cell size) in a time point, the prediction errors are tested against random choices of original parameters of all cells for all time points in the same experiment. For each test in each time point each condition in each experiment, 1,000,000 such random choices are performed. The p-values are corrected via Bonferroni-Holm correction for all time points in both experiments in the same condition.

**S1 Video. Synthesized PC12 cell differentiation in response to treatment of NGF for the first cell in Fig 5**. Cell shapes were inferred at 2.4h time steps aiming to illustrating smoother and more realistic transitions.

**S2 Video. Synthesized PC12 cell differentiation in response to treatment of NGF for the second cell in Fig 5**. Cell shapes were inferred at 2.4h time steps aiming to illustrating smoother and more realistic transitions.

**S3 Video. Synthesized PC12 cell differentiation in response to treatment of NGF for the third cell in Fig 5**. Cell shapes were inferred at 2.4h time steps aiming to illustrating smoother and more realistic transitions.

**S4 Video. Synthesized PC12 cell differentiation in response to treatment of NGF for the fourth cell in Fig 5**. Cell shapes were inferred at 2.4h time steps aiming to illustrating smoother and more realistic transitions.

**S5 Video. Synthesized probability density maps for mitochondria during PC12 cell differentiation in response to treatment of NGF for the first cell in Fig 6.** Cell shapes were inferred in 2.4h time steps to provide smoother transitions. The density is shown with a hot-cold color map similar to Fig 6.

**S6 Video. Synthesized probability density maps for mitochondria during PC12 cell differentiation in response to treatment of NGF for the second cell in Fig 6.** Cell shapes were inferred in 2.4h time steps to provide smoother transitions. The density is shown with a hot-cold color map similar to Fig 6.

**S7 Video. Synthesized probability density maps for mitochondria patterns during PC12 cell differentiation in response to treatment of NGF for the third cell in Fig 6.** Cell shapes were inferred in 2.4h time steps to provide smoother transitions. The density is shown with a hot-cold color map similar to Fig 6.

**S8 Video. Synthesized probability density maps for mitochondria patterns during PC12 cell differentiation in response to treatment of NGF for the fourth cell in Fig 6.** Cell shapes were inferred in 2.4h time steps to provide smoother transitions. The density is shown with a hot-cold color map similar to Fig 6.

